# High-throughput phenotypic profiling of insecticide responses in mosquito larvae

**DOI:** 10.64898/2026.09.16.752149

**Authors:** Kaetlyn T. Ryan, Kathy Vaccaro, Leonardo R. Nunn, Lyric C. Bartholomay, Mostafa Zamanian

## Abstract

Extensive use of insecticides is increasing resistance risks that could severely reduce the control of mosquito vectors that transmit infectious agents of Neglected Tropical Diseases. Addressing resistance threats by discovering new insecticides can be challenging because of the limited throughput and translatability of existing *in vitro* screening pipelines. Acquiring multiple phenotypic endpoints of larvae could alleviate these restrictions by more thoroughly profiling drug effects on whole organisms, providing leads to compounds with novel mechanisms of action, and increasing screening throughput compared to adult-stage assays. Here, we establish a pair of assays for profiling motility and development traits in larval-stage mosquitoes at a higher scale than standard larvicidal screening techniques. We optimized assay parameters and developed novel image processing approaches that enable relatively high throughput screening of chemical compounds on single larvae within a screen with condition replicates. We tested the assay with an insect growth regulator (S-methoprene), a slow-acting pyrrole (chlorfenapyr), and two microbial larvicides (Spinosad and *Lysinibacillus sphaericus*). These measurements aligned well with known insecticide mechanisms, and dose response curves established assay baselines for comparison in future screens. Finally, we present ways in which the assay design can be modified across different imaging technologies, showing the flexibility of the screening approach.

**Author Summary:** Insecticides are often used to control the transmission of Neglected Tropical Disease agents that are spread by mosquitoes. Growing risks of resistance necessitates assessing the efficacy of current insecticides and efforts to discover novel drugs. Performing these types of research can be challenging because of the complexity of mosquito life cycles and the limited throughput of existing screening pipelines. To address these restrictions, we developed a pair of assays that profile the motility and development of mosquito larvae after drug exposure and involve relatively high-throughput imaging and data processing. We then optimized these protocols to work with three species of mosquitoes and validated the assay by profiling the effects of four frequently used insecticides. Measurements and observations with this screen could be partially explained by the known mechanisms of these treatments, and dose response curves provided potential comparison points for future screens. We also showed the flexibility of this screening method to different imaging approaches, adding potential for use across laboratories with different resources.

## Introduction

Mosquitoes transmit the pathogens that infect billions and kill hundreds of thousands of people each year [1]. Chemical based control interventions greatly reduce mosquito-borne disease (MBD) infection transmission [2–4], but these strategies rely on only a few widely used classes of insecticides [5]. Growing resistance to these resources [6] presents an urgent need for the discovery of novel and unique active ingredients. Historically, insecticide screening has relied on labor-intensive, low-throughput, and subjective screening techniques [7,8]. However, there are several approaches that could address these limitations, and developing better screening approaches could both assess resistance to existing insecticides [9–11] and increase the likelihood of novel compound identification.

Insecticides with new mechanisms of action hold potential to mediate resistance challenges [12], and finding such compounds does not require predicting promising mosquito targets. Assessing multiple mosquito phenotypes beyond standard toxicity endpoints has previously been shown to be useful in differentiating treatment effects even if the chemicals have related putative targets [13]. Additionally, multifaceted whole-organism phenotypic screening has been successful at identifying promising active compounds and targets in other pathogen drug discovery efforts [14,15]. The throughput of mosquito screening assays can also be increased to expedite novel insecticide detection by optimizing assays to be flexible and automated in plate setup, data acquisition, or data processing. For example, assessing effects in smaller vessels than cups and cages can save large quantities of compound and space while screening [16]. Pairing imaging systems and image analysis software can both save time and yield more comprehensive phenotyping results than manual measurements [17]. Additionally, screening mosquito larvae instead of adults can increase assay throughput while still identifying pesticides that may be active against adults [18].

Designing holistic phenotyping pipelines can also help increase the understanding of known insecticides. While mechanisms of action have been established for primary insecticide classes, this does not necessarily predict how mosquitoes will react to treatments *in vitro*. Establishing metrics for these active ingredients could generate useful comparison points for future high-throughput screens. This also appears to be a useful categorization approach for projects focused on the repurposing of agrochemicals [19] or the reevaluation of insecticides never brought to market [20].

Here, we present a pair of larval phenotyping assays that attempt to address the need for higher throughput and more comprehensive screening methods. We measured both larval motility and development phenotypes and optimized these assays to be flexible across three mosquito species: *Aedes aegypti*, *Aedes albopictus*, and *Culex pipiens*. The data acquisition of these screening approaches incorporates automated imaging techniques and innovative tools for image processing. We then used these protocols to observe *in vitro* effects of four known insecticides with unique active ingredients. Finally, we explored whether this assay design could be adapted to other laboratory setups by comparing findings across different imaging systems.

## Methods

### Mosquito husbandry

*Aedes aegypti* (Liverpool strain), *Aedes albopictus* (Missouri strain), and *Culex pipiens* (Iowa strain) mosquitoes were maintained in colony at the University of Wisconsin-Madison. Larvae were reared in enamel pans and fed daily with a slurry of ground TetraMin tropical flakes (Blacksburg, VA). Adult mosquitoes were housed in an environmental chamber kept at 27°C and 80% relative humidity (RH) with a 16:8 hour (L:D) photoperiod. Adults of all species were provided with 10% sucrose solution (w/v) *ad libitum*. Once a week, females were fed a blood meal of defibrinated sheep blood (HemoStat Laboratories, Dixon, CA) provided with an artificial feeding system at 37℃ through stretched Parafilm M (Bemis Company, Inc). *Aedes* spp. eggs were collected on a damp and autoclaved seed germination paper (Anchor Paper) lining an oviposition cup placed in the colony cage for 72 hours. Papers, referred from here as egg sheets, were then removed, dried, and stored in individual Whirl-Pak bags (Nasco Sampling LLC, Madison, WI) in the environmental chamber for 2-3 months prior to assay use [21]. *Cx. pipiens* egg rafts were collected by placing an oviposition cup in the colony cage for 24 hrs. Following oviposition, eight to ten egg rafts were transferred to mason jars with diH_2_O for assay preparation [9].

### Motility and development assay set up

*Aedes* spp. eggs were hatched synchronously by submerging egg sheets in 100-500mL of autoclaved, deoxygenated water in sterile glass mason jars stored at 27°C for 1-2 hours. *Cx. pipiens* eggs were asynchronously hatched at 27°C across 16-24 hours by transferring egg rafts from the colony egg dish to a sterile glass mason jar containing 500mL of distilled water supplied with a small quantity of finely ground and diluted TetraMin® tropical fish flakes (Spectrum Pet Brands LLC, Blacksburg, VA), referred to as fish food. Hatched larvae were poured into clear glass dishes for easy pipetting, and one or five first instars were transferred via pipette to individual wells of a 96-well plate (Greiner Bio-One Cat. No 655-180) in aliquots of 50µL. Plates with five larvae per well were used for acquiring motility data and plates with one larva per well were used for acquiring motility and development data.

Next, volumes of distilled water were added depending on the final treatment plan for the well: 50µL for treatments or controls in water solvent (Vectolex, Spinosad, Altosid) and 100µL for treatments or controls with DMSO solvents (Chlorfenapyr). Food slurry was prepared by combining 0.33g of finely ground TetraMin Tropical Flakes fish food and 50mL diH_2_O for 5 larvae/well plates. This slurry was diluted 2:3 (v:v) in diH_2_O for 1 larva/well plates. Next, the food slurry was mixed thoroughly, transferred to a trough, and then added to plate wells in 50µL aliquots using a multichannel pipette, mixing the solution between each row. Plates were sealed with breathable strips (Diversified Biotech BERM-2000) and placed in humidity chambers prepared as described previously [22] that were stored in a shaking incubator at 28°C and 30rpm.

### Drug preparation and addition

VectoLex WSP (Valent BioSciences LLC, Libertyville IL) powder was used for *Lysinibacillus sphaericus* treatment and dissolved in diH_2_O immediately prior to any assay use with thorough vortexing. Spinosad (Natular G30, Clarke Mosquito Control Products Inc, St. Charles IL) powder and Altosid (Zoëcon, Wellmark International, Schaumburg IL) solution (used for S-methoprene treatment) were diluted in diH_2_O as assay aliquots that were stored at room temperature. Chlorfenapyr (CAS #122453-73-0 Cayman Chemical Company, Ann Arbor MI) powder was stored at -20°C and dissolved in DMSO to make assay aliquots that were similarly stored. Ivermectin (CAS #70288-86-7 Fisher) powder was stored at 4°C and dissolved in DMSO to make assay aliquots. Water-based treatments were made as 4X aliquots and DMSO-based treatments were made as 100X aliquots.

Larval plates were incubated post-food addition for a timeframe specified to the species and number of larvae per well prior to drug addition (0 hours for *Cx. pipiens* 1/well plates, 24 hours for *Cx. pipiens* 5/well and *Aedes spp*. 1/well plates, and 48 hours for *Aedes* spp. 5/well plates). See Results for a description of how these metrics were determined. After this incubation period, drug aliquots were added to wells at quantities of 50µL for water-based treatments or 2µL for DMSO-based treatments. Plates were then resealed with breathable strips (Thermo Scientific) and returned to humidity chambers inside the shaking incubator until and between imaging timepoints described below.

### Motility image acquisition

Plates were imaged 1, 24, 48, and 72 hours post-treatment. For *Aedes* spp. plate readings on the ImageXpress Nano (Molecular Devices), images were collected by well with a 2x objective for 10 consecutive frames (3.3 frames per second). For *Cx. pipiens*, low baseline motility rates were noted, so videos acquired on the ImageXpress were taken with staggered frames by taking an image of each well of the plate and repeating this process 9 times. For plate readings on the 6-camera Loopbio Motif system, recording durations of 30 seconds, 2 minutes, and 5 minutes were used. The recording codec used was hq-fast at a max framerate of 15. All motility readings were performed at room temperature.

### Development (size) image acquisition

After 72 hour motility image acquisitions, plates were frozen at -20°C for 24 hours. Frozen 1 larva/well development plates were left at room temperature with lid removed for 24 hours to allow plates to thaw and water to evaporate until 50µL of water or less remained in the wells to get larvae close to the bottom of the well (the focus range of still images) without damaging their tissue. Next, 75µL of vegetable oil was added to each well while using pipette tips to push any larvae stuck to the edges of wells to the plate bottom during the process. Plates were then imaged using the ImageXpress Nano (Molecular Devices) with a 2x objective for a single frame. Plates containing 5 larvae/well plates were disposed of after freezing.

### Image and data analysis

Mosquito motility videos were processed to generate optical flow values using the motility module of wrmXpress v1.4.0 [23] (docker image v6) through a node maintained by UW-Madison’s Center for High Throughput Computing. The instance segmentation model used for processing development images was trained using Roboflow, YOLOv8, and Ultralytics softwares. For training, 672 images across varying timepoints, mosquito species, and food quantities were annotated and augmented using flip, 90° rotation, and blur steps. The image dataset was then split for training (87%), validation (8%), and testing (4%). YOLO training was performed with 100 epochs and a standard resizing value of 640. The model (mosquito_larvae_segment2.pt) was added to wrmXpress pipelines, and wrmXpress v2.0 was used to process development plates. Motility and development output data was analyzed using R software including the tidyverse and drc packages. Motility and development values were normalized as follows: (X - positive control) / (negative control - positive control) where X is the phenotypic endpoint value.

## Results

### Optimizing assay growth conditions and treatment timing across mosquito species

To acquire motility and development endpoints, we decided to use optical flow (the pattern of object movement within a sequence of images) and larva size quantified by pixel count as respective proxies. Establishing baselines and benchmarks of these phenotypes required optimizing several assay conditions across mosquito species, including the density of larvae in individual wells, the amount of food provided, and acclimation timing. Plates prepared with 5 larvae per well showed more consistent motility readings across replicates because singular larvae were at times not visible due to well shadowing or occlusion by food. However, when 5 larvae occupied a well, competition for food led to asynchronous larval development, obscuring treatment effects on size. To address these complications, plates containing 5 larvae/well were measured for motility effects only, while optical flow rates and larva size were measured in plates with 1 larva/well to determine if development effects are independent of motility effects.

A single aliquot of food was used for assays to avoid diluting drug treatments. Because of this, we had to determine what food quantities were sufficient to feed larvae through development stages without visually obscuring them during imaging (**Fig 1A**). Ultimately, plates were initially supplemented with 300µg of fish food for 5 larvae per well plates and 120µg for 1 larvae per well plates, and these quantities functioned well across all species of mosquito tested. However, food consumption and development varied across well densities and species. To mediate this, we evaluated various acclimation periods and treatment timings to identify the ideal imaging window for each species and density pair (**S1A and S1B Figs**). We aimed to ensure larvae had consumed enough food for early motility detection but were young enough to allow for longitudinal development analysis.

**Fig 1.**
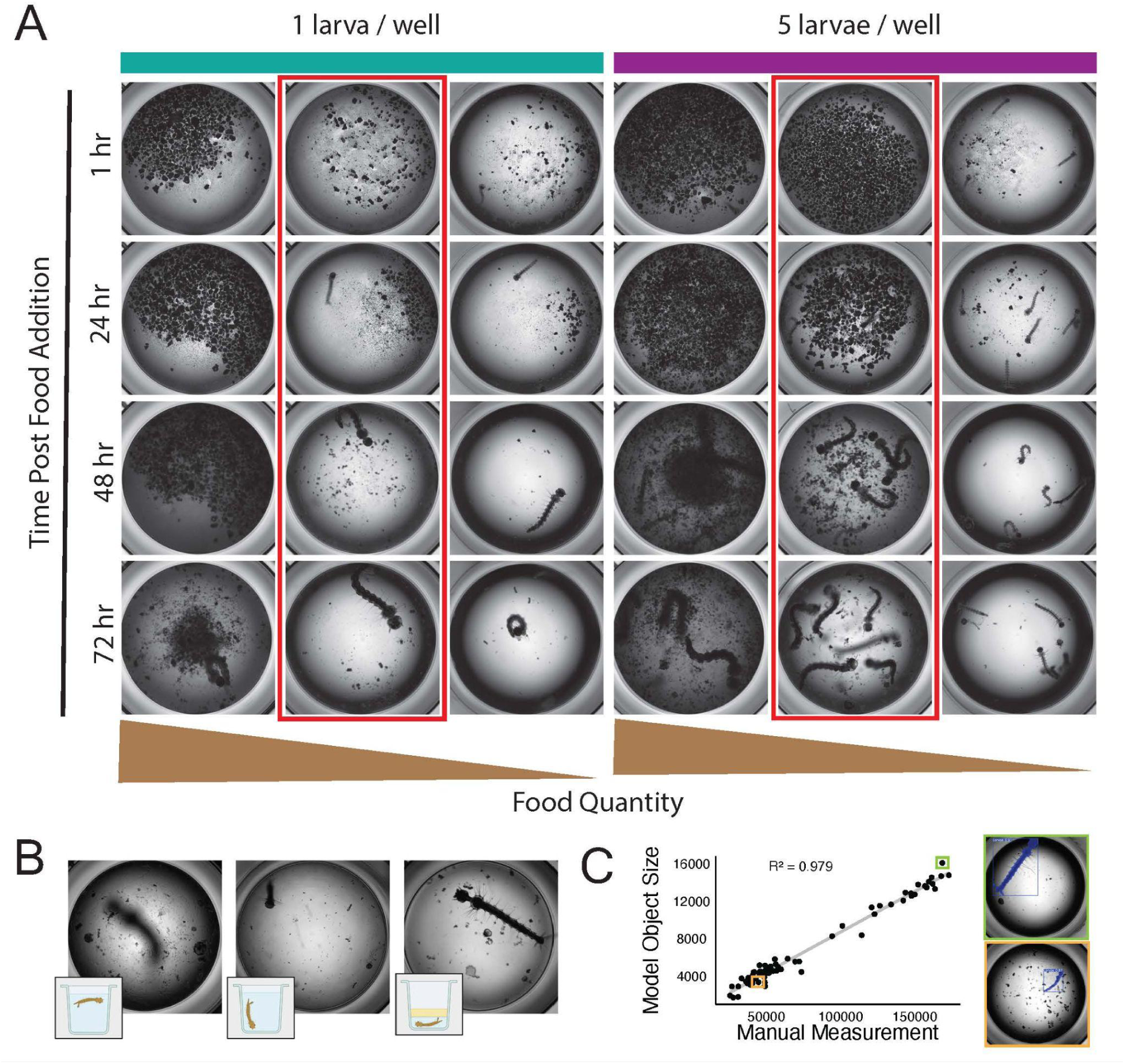
Optimized parameters of motility and development assays. (A) Representative images of larval growth patterns across time after different initial feeding quantities in plates with 1 or 5 larvae per well. (B) Representative development images of floating larvae (left), vertical larvae (middle) or larvae flattened by a film of vegetable oil (right). (C) Comparison of mosquito sizes after processing images manually (Fiji) or with the developed object detection model (YOLOv8) and examples of large (green) and small (orange) larvae images post model processing.

*Cx. pipiens* larvae had significantly lower baseline motility rates than *Aedes* larvae, so different imaging techniques had to be tested to distinguish between motile and non-motile *Cx. pipiens*. We attempted to provoke movement by using the shutter of the ImageXpress to flash transmitted light or by exposing larvae to fluorescent wavelengths, but the highest baseline motility rate was achieved when frames were collected across the entire plate instead of sequentially per well, likely because this increased the likelihood that larvae would be in different positions between frames (**S1C Fig**). To accurately measure larval size, plates were frozen and liquid was evaporated from wells before a layer of vegetable oil was used to acquire clear images of flattened larval bodies in a focused 2D frame (**Fig 1B**). To increase the efficiency of measuring larvae size, a YOLOv8 [24] model was trained to detect and box larvae within images. The model exhibited precise and thorough object detection when tested with annotated images (**S2 Fig**), and size measurements aligned closely with measurements manually collected using Fiji [25] software (**Fig 1C**).

### Detecting larvicidal effects of known insecticides

Assay parameters selected for each species and well density are summarized in **Fig 2A**. These protocols were used to assess the motility and development effects of four commercial insecticides: Altosid, chlorfenapyr, Spinosad, and VectoLex. In these screenings, water or DMSO solvents were used as negative controls and 10 µM ivermectin was used as a positive control because of its known lethal effect on larvae [26–28]. Motility and development readings of treated larvae were normalized between these two control values and displayed as dose-response curves in **Fig 2B** and **Fig 2C**. Calculated EC50 values for each treatment across species and timepoints are summarized in **S1 Table** and **S2 Table**.

**Fig 2.**
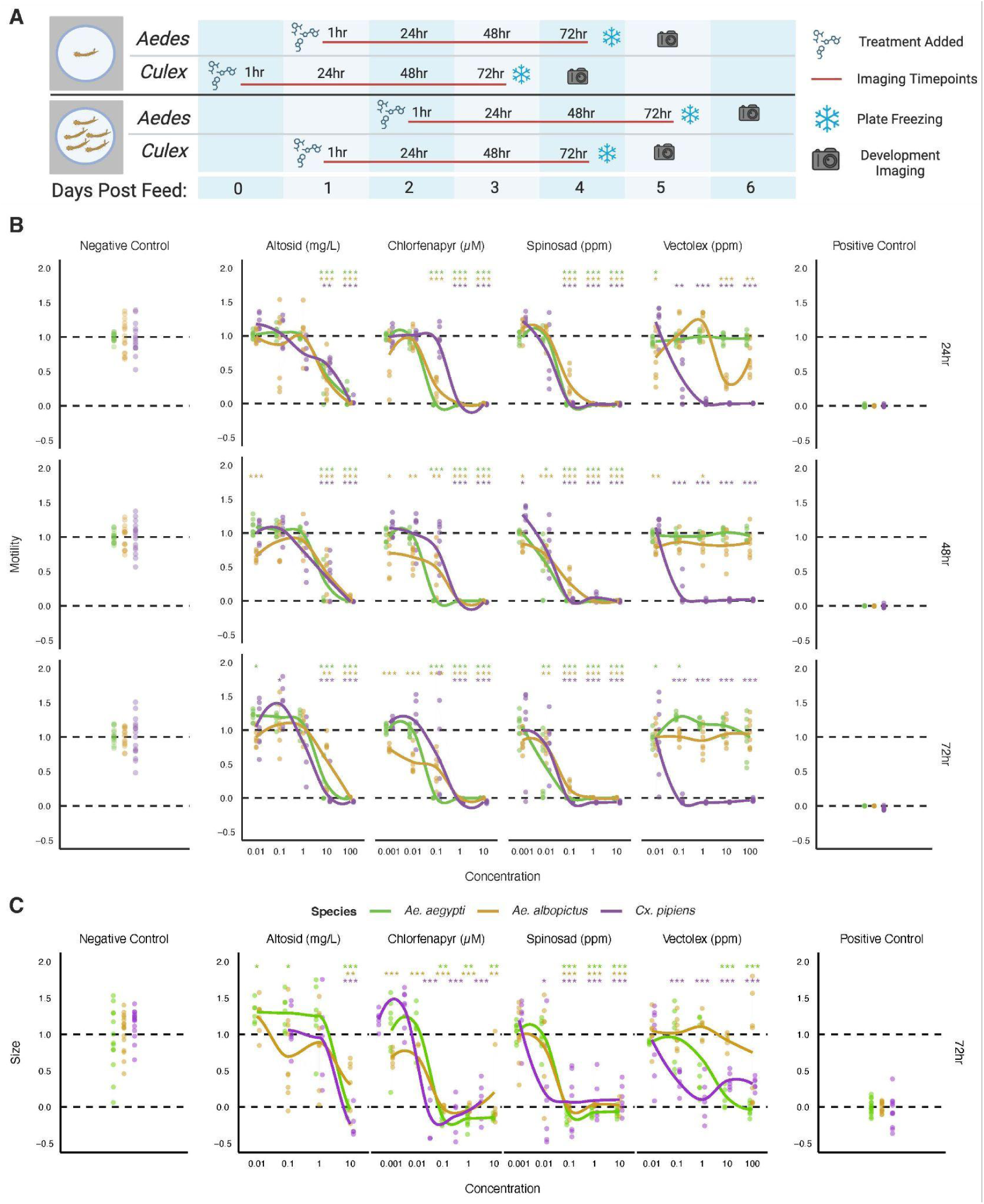
Insecticide findings of motility and development screening. (A) Schematic summarizing the treatment timing and data collection for assays across mosquito species. (B) Normalized motility measurements of three mosquito species (color) after insecticide treatment across varying concentrations (x-axis). Data from the 1 hour timepoint is not shown. (C) Normalized mosquito size measurements of three mosquito species (color) after insecticide treatment across varying concentrations (x-axis). Statistical analyses were performed via T-test and reported as follows, * : p<0.05, ** : p<0.01, *** : p<0.001, **** : p<0.0001.

Altosid only had motility and development effects at high concentrations (≥10mg/L), and EC50 values were slightly lower for development measurements. Chlorphenapyr motility effects were more obvious over time in *Aedes* species, and development curves were very similar to 1 larva per well motility curves (**S3 Fig** and **S3 Table**). Spinosad motility and development results were similarly aligned and consistent across species and timepoints. VectoLex had stronger effects on motility and development of *Cx. pipiens* larvae than *Aedes* species.

### Assay adaptability to alternative imaging systems

Most of the data acquisition for this study involved using a high-content imaging device (ImageXpress, Molecular Devices) that records wells individually and sequentially. However, we also investigated how the assay conditions would perform with a different imaging system using a 6-camera Loopbio instrument that images plate wells simultaneously in groups of 16 from above (**Fig 3A**). Plates of *Ae. aegypti* larvae treated with a range of chlorfenapyr, Spinosad, and ivermectin concentrations were prepared to collect a breadth of motility rates before being acquired by both imaging systems (**Fig 3B**). All imaging data was processed using the wrmXpress [23] optical flow algorithm and compared using a linear regression model. Short Loopbio video recordings (30 seconds) did not align as well with ImageXpress readings as 2 minute recordings did (R^2^ values of 0.50 and 0.74 respectively), but R^2^ values of approximately 0.74 were consistent with longer (5 minute) Loopbio recording times.

**Fig 3.**
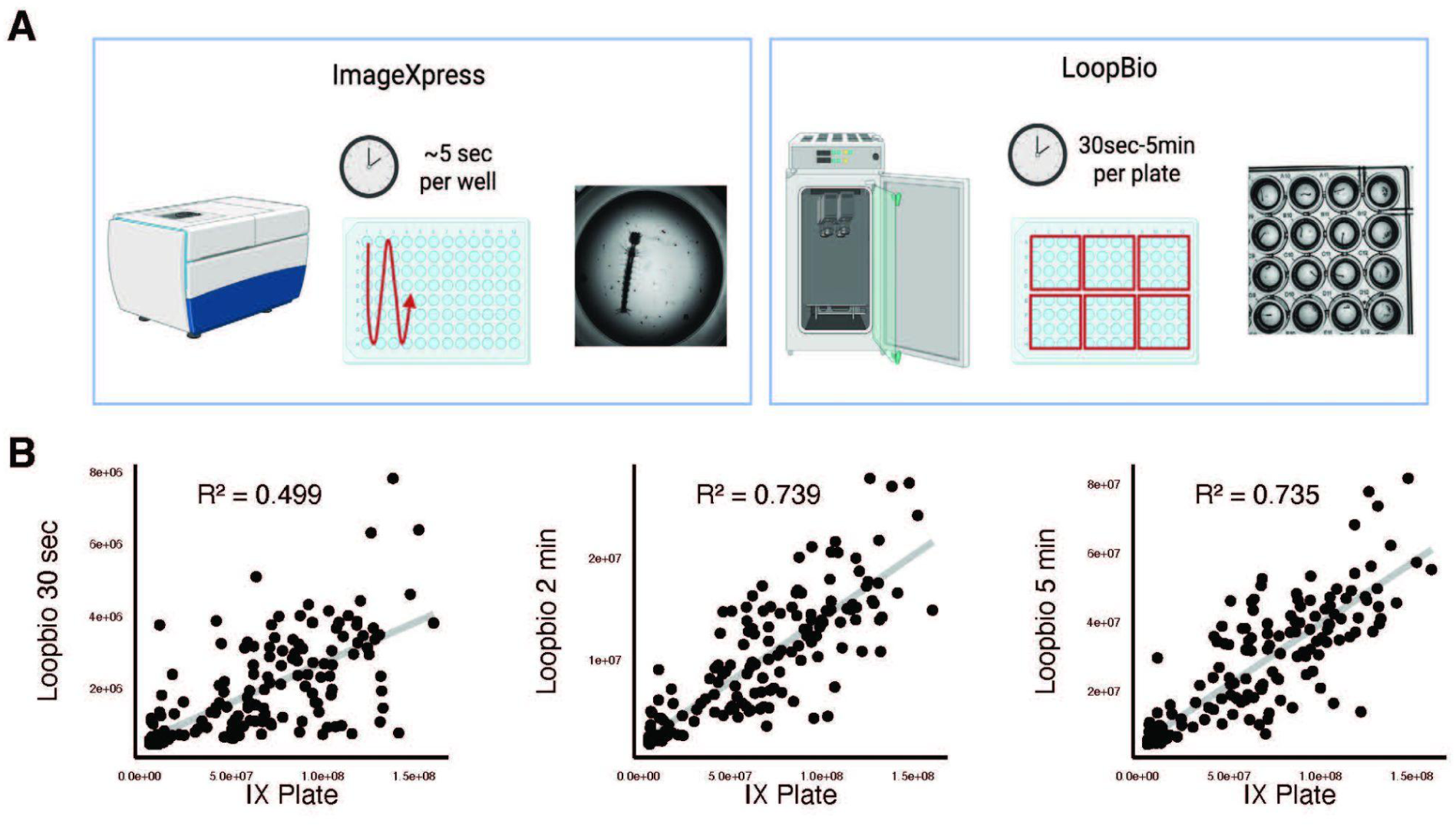
Comparison of motility assay results across two different imaging systems. (A) Schematic showing time periods, imaging order, and image output examples of ImageXpress (left) and Loopbio (right) imaging systems. (B) Comparisons of plate measurements between ImageXpress (IX) and measurements recorded on the Loopbio system over 30 seconds, 2 minutes, and 5 minutes.

## Discussion

In this study, we performed the stringent optimization required to design large-scale, multivariate phenotyping assays for larvae, and applied these screening techniques to describe *in vitro* effects of known insecticides. Some benefits of these methods resulted from the ways we aimed to increase assay throughput compared to historical screening approaches. For example, standard screening protocols utilize full cups for each treatment condition [7], but our plate-based approach uses volumes of 200µL per technical replicate, greatly reducing the amount of compound material and laboratory space needed for assay completion. Additionally, many established screening techniques involve probing and observing individual larvae or using alternative methods of manual movement stimulation [7,8], but by exploring precise imaging techniques, we showed that these processes can be expedited by automating measurement and quantification (**S1C Fig**). The remaining bottleneck of our assay throughput is in how larvae are manually distributed to wells. This is the most time consuming step of our methods, but the process could likely be automated with large particle liquid handling instruments that are already used in transgenic studies [29].

Another benefit of this screening method is the ability to detect more cryptic phenotypes than previous protocols that focus exclusively on larvae death rates. Some labs and organizations have successfully designed larval assays with high throughput levels, but predicting *in vivo* effects from *in vitro* results can still be challenging [30]. Looking at a breadth of phenotypes could reduce rates of false negatives in screening and provide early leads for modes of action studied in follow up [13]. Precise image analysis also generates more quantitative results that can better discern between small effects like those studied in early resistance assessments [9,31]. One reason phenotypic imaging strategies like these have not been widely integrated yet is likely because of the many assay parameters that must be optimized and the variation in responses across mosquito species as we describe (**Fig 1 and S1 Fig**). Feeding quantities and timing are examples of components that we carefully parsed but could also be further refined when considering impacts on drug intake, especially when studying insecticides that act through consumption.

Using this methodology, we observed phenotypic outcomes for mosquito larvae that aligned with the mode of action for the active ingredient of each insecticide tested (**Fig2**). For example, Altosid had effects on both larval motility and development at similar concentrations (≥10mg/L). This is consistent with findings that the active ingredient (S-methoprene) and related compounds inhibit larval molting [32]. Because treatments were applied to early instars, low motility rates could indicate either paralysis or lower optical flow rates of developmentally inhibited larvae compared to full grown controls. The fact that a phenotypic response to Altosid was only detected at high concentrations could be explained by previous findings that late instar larvae stages are more sensitive [33,34] than the early stages at which treatment was applied in the assay. Chlorfenapyr requires P450 activation to produce biotoxins [35], and this could explain our observations that chlorfenapyr motility inhibition at lower concentrations increases at later timepoints. Additionally, development curves are similar to the 24 and 48 hour motility curves implying that early paralysis is responsible for inhibited development. Spinosad and VectoLex are also known to be lethal to mosquito larvae [36,37], and screening results here show motility and developmental stunting occur at similar concentrations and timepoints. This study does not show an example of a treatment that influences motility or development effects independently, but pairing these two phenotypes could show such a result in a future screen and lead to a novel insecticide with a unique mechanism of action.

EC50 values could vary across species in part because of the differences in treatment timing across assay set ups. For example, *Cx. pipiens* larvae generally had lower VectoLex EC50 values, and this may be because these larvae have less acclimation time in the plate than *Aedes* species do, not because they are more susceptible to treatments. Alternatively, this variation may be attributed to the slightly advanced stage of the *Cx. pipiens* and pre-plating feeding this species receives as described in the methods. These factors could cause larvae to ingest VectoLex faster and lead to activity at lower concentrations because the substance becomes lethal in the gut lining.

Strong motility correlations were achieved (R^2^ values of ∼0.74) across different imaging devices, and remaining discrepancies can be explained by some of the advantages and limitations of each system. When acquiring motility data with the ImageXpress, the larvae have to be captured within a focus range near the plate bottom for optical flow to be detected, so larvae swimming at the top of the well are not recorded. Alternatively, measuring whole plates with the Loopbio system captures larvae at all well depths but often results in more shadowing within wells and lower resolution. In the comparison presented in **Fig 3**, only 1 larva/well plates at the 24 hour timepoint were compared, so these correlations may also vary across different species, well densities, and timepoints. However, both imaging systems detect a wide range of motility values, indicating that the assay design is flexible across equipment types.

The fact that we were able to describe multiple phenotypic responses of known insecticides at a relatively high throughput indicates that this assay would be useful for both novel insecticide screening and resistance monitoring across mosquito populations [9]. Because the assay acquires phenotypes of individual larvae, it could also be applied to other projects like screening for marker expression in transgenic mosquitoes [38] or measuring effects of entomopathogens on mosquito development and survival [39,40]. The assay could also be further adapted to capture other phenotypes or be extended to show development through pupation stages. The integration of the larvae segmentation model into the open-access wrmXpress software [41] and flexibility of the assay across devices makes the method accessible to any lab with an imaging system suited for 96-well plates. In conclusion, we have established and validated a high throughput assay for measuring larval motility and development responses that could be adapted for a diverse array of purposes in the study of vector biology and mosquito control.

## Acknowledgements

This work was supported by National Institutes of Health NIAID grants R01 AI151171 to M.Z. Thank you to Ali Ross for providing insecticide resources. Thank you to members of the Midwest Center of Excellence for Vector-Borne Disease for providing manuscript feedback.

## Author Contributions

KTR: Conceptualization, Data Curation, Formal Analysis, Investigation, Methodology, Writing (Original Draft Preparation)

KV: Conceptualization, Methodology, Writing (Review & Editing)

LRN: Software, Writing (Review & Editing)

LCB: Conceptualization, Resources, Validation, Writing (Review & Editing)

MZ: Conceptualization, Funding Acquisition, Project Administration, Supervision, Writing (Review & Editing)

## Supporting Information

**S1 Fig. Additional motility assay optimization components.** (A) Representative images showing *Aedes* spp. growth patterns over 72 hours (x-axis) after 24 or 48 hours of plate acclimation (y-axis) prior to control treatment. (B) Representative images showing *Culex pipiens* growth patterns over 72 hours (x-axis) after 0, 24, or 48 hours of plate acclimation (y-axis) prior to control treatment. Red boxes indicate procedures chosen for the assay. (C) Different approaches attempted to capture *Cx. pipiens* motility with ImageXpress settings including the settings used for *Aedes spp.* plates (Protocol), opening and closing the shutter to induce light exposure (Shutter Flutter), imaging over a longer period (20 Frames), exposing larvae to GFP, Texas Red, and DAPI wave lengths (WL) prior to transmitted light imaging, and imaging each well of the plate and repeating to capture separated frames (Cross Plate).

**S1 Table. EC50 values of 5 larvae/well motility screen.** NA values indicate conditions that either had no effects or warnings during calculations.

**S2 Table. EC50 values of development screen.** NA values indicate conditions that either had no effects or warnings during calculations.

**S2 Fig. Quality control measurements of the object detection model.** (A) Comparison of precision and confidence metrics for tested object boxes. (B) Comparison of recall and confidence metrics for tested object boxes. (C) Normalized confusion matrix showing the proportion of tested images that resulted in false positives, false negatives, true positives, and true negatives. (D) Precision, recall, mean average precision at intersection over union threshold of 50, and mean average precision at varying intersection over union threshold between 50 and 95 across training epochs.

**S3 Fig. Normalized motility values for 1 larva/well plates.** Data from the 1 hour timepoint is not shown. Statistical analyses were performed via T-test and reported as follows, * : p<0.05, ** : p<0.01, *** : p<0.001, **** : p<0.0001.

**S3 Table. EC50 values of 1 larva/well motility screen.** NA values indicate conditions that either had no effects or warnings during calculations.

