## Supplementary figures and images for "High-throughput phenotypic profiling of insecticide responses in mosquito larvae"

### S1Fig

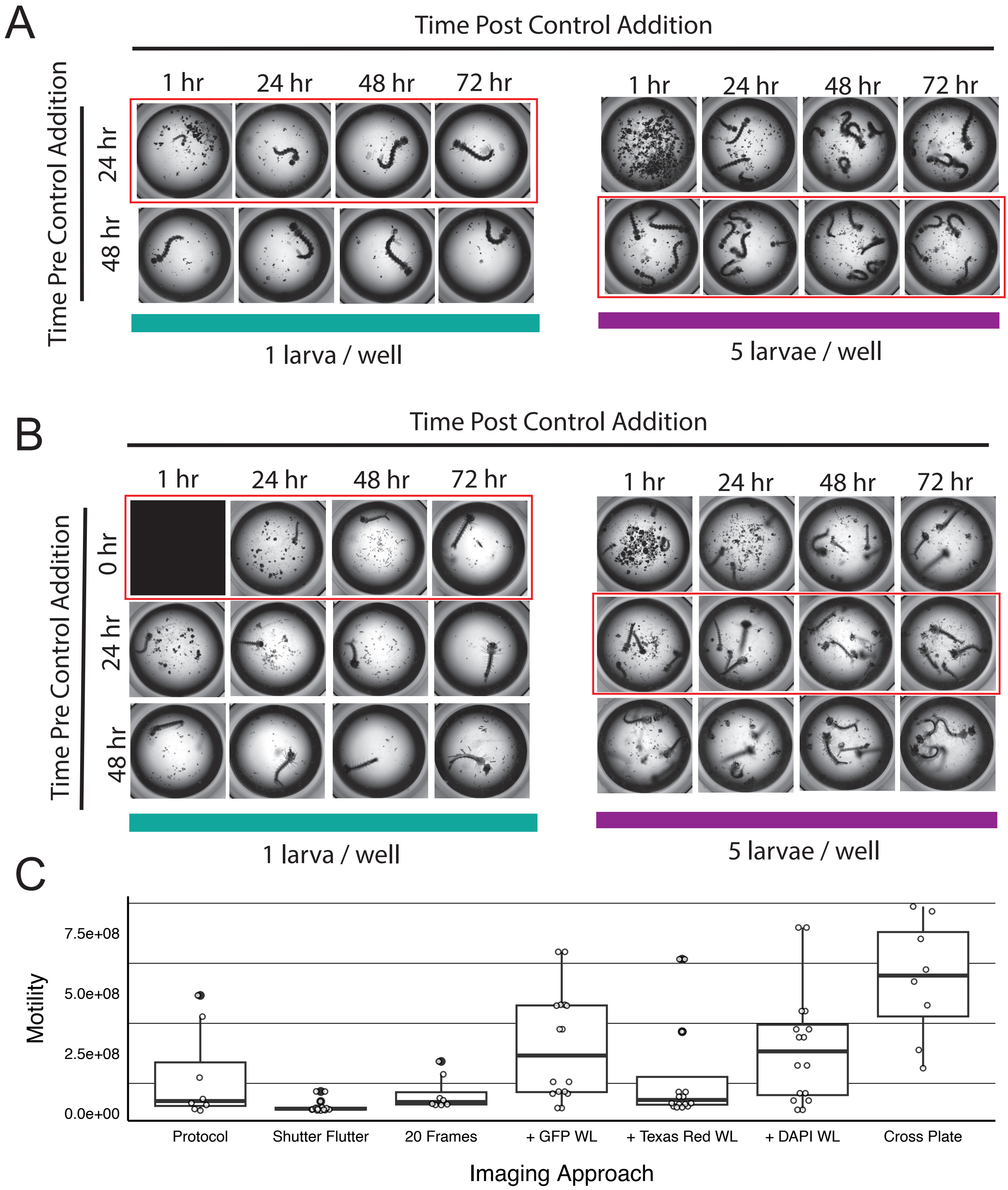

### S2Fig

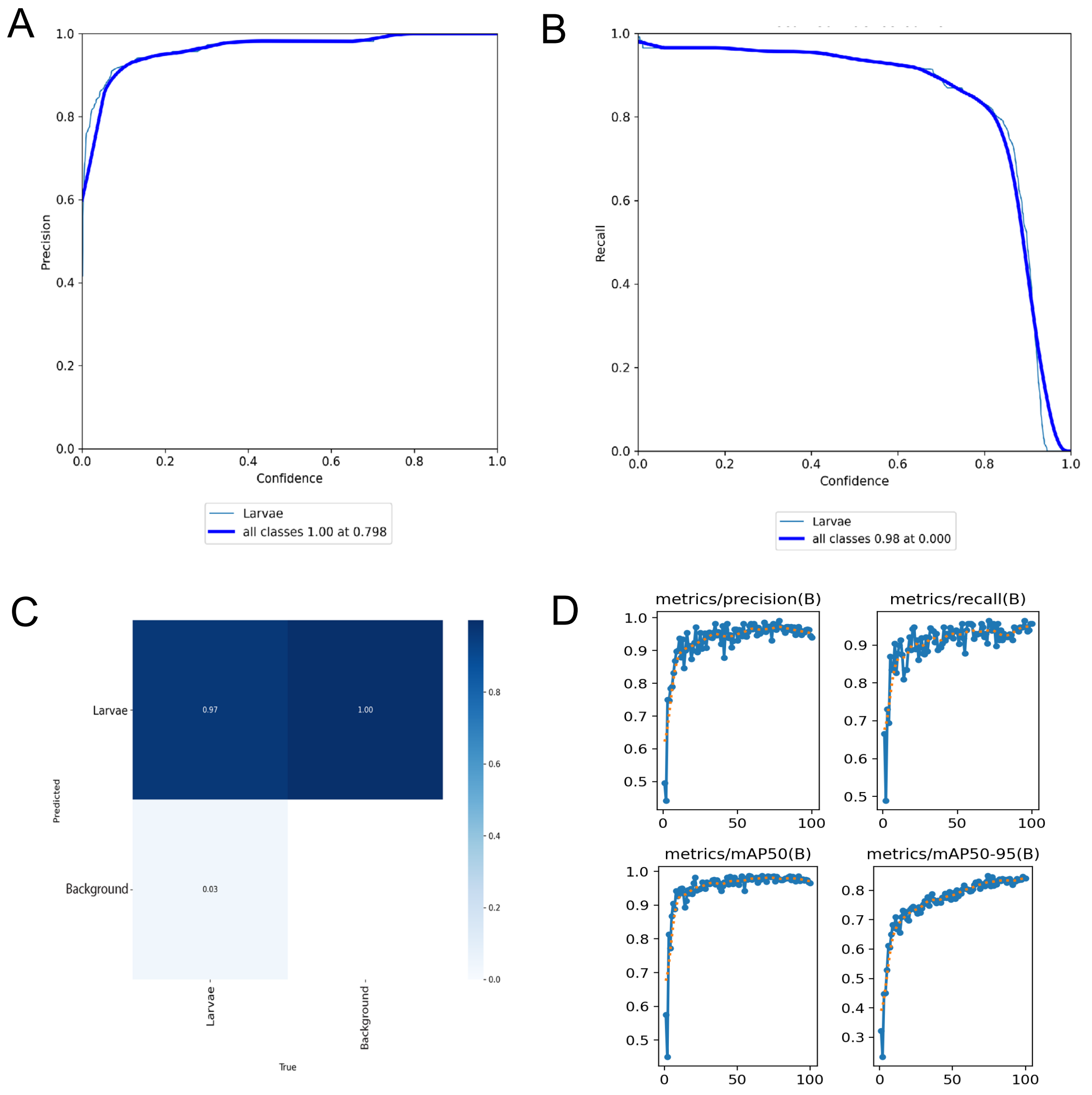

### S3Fig

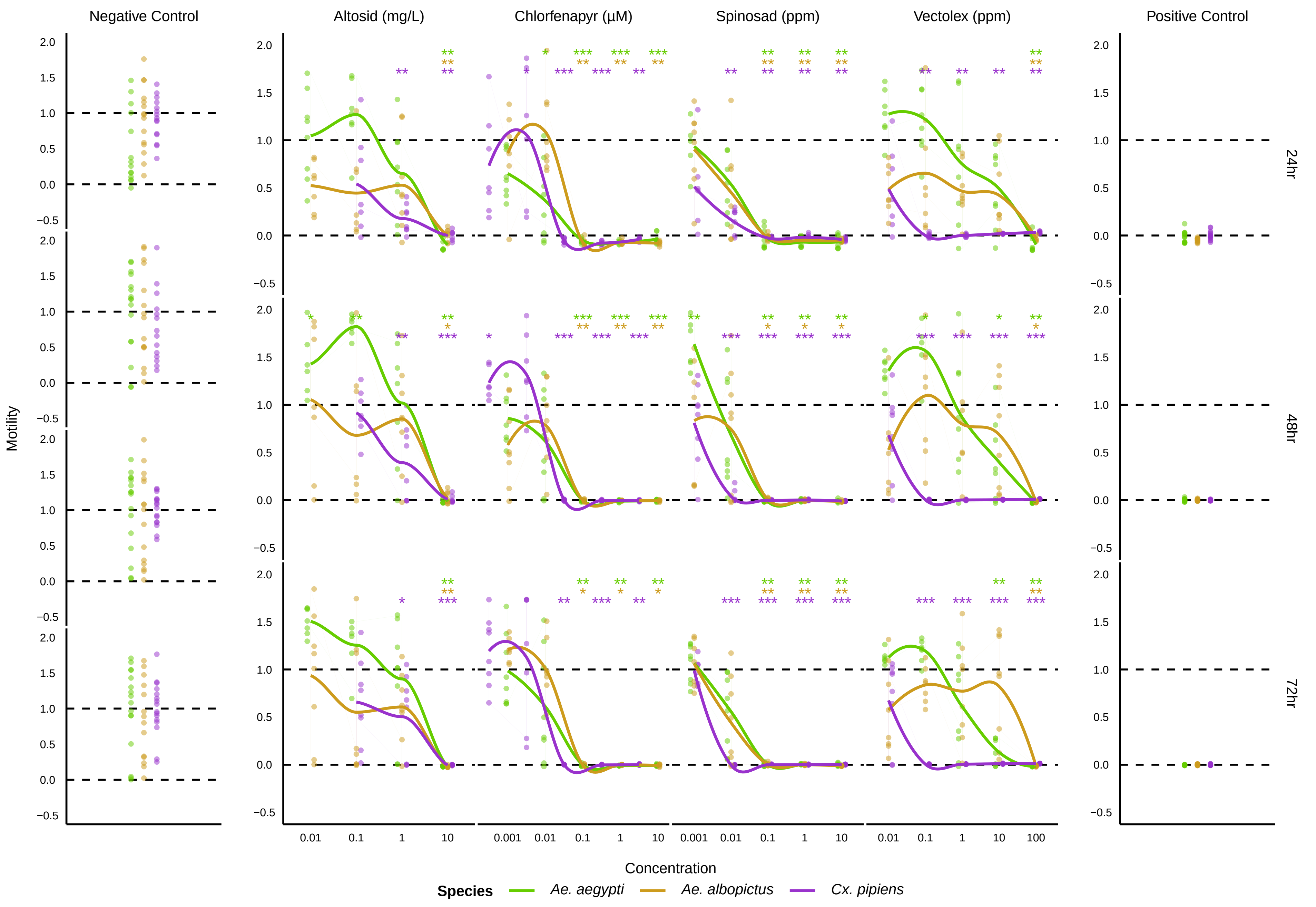
